# Targeted RNA knockdown by crRNA guided Csm in zebrafish

**DOI:** 10.1101/228775

**Authors:** Thomas Fricke, Gintautas Tamulaitis, Dalia Smalakyte, Michal Pastor, Agnieszka Kolano, Virginijus Siksnys, Matthias Bochtler

**Affiliations:** International Institute of Molecular and Cell Biology, Trojdena 4, 02-109 Warsaw, Poland; Institute of Biotechnology, Vilnius University, Saulėtekio av. 7, 10257 Vilnius, Lithuania; Polish Academy of Sciences, Institute of Biochemistry and Biophysics, Pawinskiego 5a, 02-106 Warsaw, Poland

## Abstract

RNA interference (RNAi) is a powerful experimental tool for RNA knockdown, but not all organisms are amenable. Here, we provide a “proof of principle” demonstration that CRISPR endoribonucleases can be used for programmable mRNA transcript degradation. Using zebrafish as the animal model and Csm(crRNA) complexes as the CRISPR endoribonucleases, we have targeted a transgenic EGFP transcript expressed from a variety of promoters. A drastic decrease of fluorescence was achieved in germ cells of the vasa:EGFP line. Weaker effects were also seen in fish lines that express EGFP zygotically. Knockdown was statistically significant in cmcl2:EGFP and fli1:EGFP zebrafish lines at 1 day post fertilization (dpf), but reduced to background levels at 2 dpf. The nkx2.5:EGFP fish line was least susceptible to Csm mediated EGFP knockdown. We conclude that at the present stage, Csm mediated knockdown is already efficient for maternal transcripts, and may compare favorably with morpholinos for such targets in zebrafish.

## INTRODUCTION

Chronic knock-out and acute knockdown often lead to drastically different phenotypes, in ways that are only partly explained by technical limitations (1,2). Therefore, knockdown experiments remain interesting even in the era of facile DNA knockout generation using CRISPR Cas9.

In many experimental animal models, systemic (3) or cell-autonomous (4) RNA knockdown can be achieved exploiting endogenous RNA inference pathways, but there are important exceptions, zebrafish among them. In zebrafish embryos, esiRNA has been reported to lead to non-specific developmental defects (5), presumably due to overload of the endogenous interference pathways. More encouraging results have been obtained using either siRNA (5) or shRNA (6), but neither method has gained widespread acceptance in the zebrafish community.

Morpholino mediated knockdown is the leading technology for RNA silencing in zebrafish (7-9). The method is well established, but has some limitations. As morpholinos only bind to RNA but do not cleave it, only translation initiation and splice sites can be targeted, which is not always uniquely possible. Moreover, many maternal RNAs are already spliced and therefore very difficult to target. Morpholinos have a low “useful” range of concentrations. At lower concentration, silencing is frequently incomplete (10). At higher concentration, off-target effects are common and have to be carefully controlled (1,11). Very recently, dCas9 mediated transcriptional inhibition (12) has been tested in zebrafish as an alternative to morpholinos, but success has so far only been demonstrated in one case in relatively early embryos (13). For maternal transcripts, morpholino-based knockdown is problematic, and transcription suppression can altogether not be used.

Prokaryotic argonaute proteins (Agos) provide an attractive, RNA degradation based strategy for programmable, targeted knockdown. The approach becomes possible because some prokaryotic Agos, such as the *Marinitoga piezophila* Ago (MpAgo), differ in their guide nucleic acid requirements from endogenous Agos (14). *Natronobacterium gregoryi* Ago (NgAgo), now considered as a DNA guided, RNA directed endonuclease (15), is to our knowledge the only heterologous Ago that has been tested as knockdown tool for zebrafish. In principle, NgAgo should make it possible to target maternal transcript as well or better than zygotic transcripts. However, to our knowledge, only knockdown of one early zygotic gene (*fabp11a*) has been tested, leading to eye development defects (15).

RNA directed CRISPR endoribonucleases, such the class1 (multiple subunits) enzymes of the Csm (type III-A) (16,17) and Cmr (type III-B) (18,19), or class 2 (single subunit) effectors such as Cas13a (C2c2, type VI-A) (20-22) and Cas13b (type VI-B) (23), or Cas9 (type II) redirected to RNA using PAMmers (24) could potentially be used for programmable RNA degradation. In this work, we have tested this concept using zebrafish as the animal model and type III-A Csm complexes as the programmable CRISPR RNA endoribonucleases.

The Csm proteins and bound crRNA form multi-subunit protein-RNA complexes (25). Initial genetic data suggested that these complexes act as RNA-guided DNA endonucleases (26). Subsequent in *in vitro* studies, however, indicated that the Csm complexes had RNA-guided, RNA-directed endoribonuclease activities cleaving substrate RNAs at multiple, regularly spaced sites (16,17). The apparent inconsistency between *in vivo* and *in vitro* data was resolved by the demonstration that the Csm complexes have transcription dependent DNase activity (27). Csm complexes find their targets by hybridization of the guide crRNA with the nascent RNA from transcription (28). They carry out co-transcriptional DNA and RNA cleavage during the expression phase of bacterial immunity, thus cleaving DNA and eliminating RNA of the invader (28,29).

Csm active sites responsible for the DNase and RNase activities are distinct and located in different subunits of the protein complex. Cas10, an HD nuclease and one of the subunits present in only a single copy in the complex, harbors the DNase activity (28-30). Csm3 subunits, present in multiple copies and forming a crRNA binding filament structure, harbor the RNase activities of the Csm complex (17). Consistent with this assignment, the Cas10 D16A and Csm3 D33A mutations (*Streptococcus thermophilus* numbering) specifically abolish ssDNase and RNase activities of the complex, respectively (17,28). Csm complexes can so far only be assembled in bacterial cells. In the natural hosts, pre-crRNAs are expressed from CRISPR region and then processed by cleavage in positions −8 and −9 of the repeat region (numbering with respect to the spacer region) and optional trimming on the 3’-side of the repeat, in steps of six nucleotides as determined by Csm3 as a ruler protein and ribonuclease (31,32). The crRNA maturation and the loading on Csm complexes also occur in heterologous hosts carrying the Cas/Csm operon and associated CRISPR region (17).

For our knockdown experiments, we used zebrafish lines expressing EGFP from various promoters in different tissues and at different stages of development. The vasa:EGFP (33), mito:EGFP (34), nkx2.5:EGFP (35), cmcl2:EGFP (36), and fli1:EGFP (37) test cases were chosen to include cases of maternal RNA deposition without zygotic expression, maternal and zygotic expression, and zygotic expression only. In all cases, the EGFP transgenes were expressed from transposon insertion sites, always in the background of the endogenous genes.

The vasa:EGFP fish have maternally deposited EGFP mRNA transcripts, and also protein. Genetic experiments show that vasa:EGFP mRNA in germ cells is exempt from the widespread RNA degradation at the mid-blastula transition and stable at least until 50 hours post fertilization (hpf). During this period, zygotic EGFP transcription is not detectable (33). Thus, the vasa:EGFP fish are good test system for knockdown of maternal RNA without confounding effects of embryonic transcription.

The mito:EGFP fish express EGFP with N-terminal mitochondrial localization signal derived from subunit VIII of cytochrome oxidase under the control of the constitutive elongation factor 1α (EF-1α) (34). Bright mitochondrial fluorescence of unfertilized oocytes suggests that maternal deposition of EGFP in mitochondria and likely also maternal mRNA deposition. As mito:EGFP fish also express EGFP in the zygote, this fish line represents a case of both maternal and zygotic transcripts (34).

The nkx2.5:EGFP fish first exhibit fluorescence in the ventral margin of the embryo at the onset of gastrulation (∼5.5 hpf). By 2 dpf, EGFP expression is limited to the heart (35). cmcl2:EGFP expression becomes detectable at ∼ 16 hpf in myocardial cells of the heart and persists for the lifetime of the animal (36). The fli1: EGFP fish start to express EGFP from the three-somite stage (∼10 hpf). At 1 dpf, trunk and segmental vessels and cells of erythroid morphology are fluorescently labelled. At 2 dpf, there is also fluorescence in the neural crest derived aortic arches and the developing jaw (37).

Therefore the hearts of nkx2.5:EGFP and cmcl2:EGFP and vessels of the fli1: EGFP reporter lines are good test cases to check knockdown of zygotically expressed EGFP in the embryo.

## MATERIAL AND METHODS

### EGFP and jGFP sequences

The target regions of the mRNAs of jellyfish (*Aequorea victoria*) jGFP and zebrafish EGFP used in this study span nucleotides 241-276. In this region, the coding strands have the DNA sequences 5’-CATGACTTTTTCAAGAGTGCCATGCCCGAAGGTTAT-3’ and 5’-CACGACTTCTTCAAGTCCGCCATGCCCGAAGGCTAC-3’ differing in seven positions (underlined).

### CRISPR regions and crRNAs

Artificial CRISPR loci were created that contained four identical 36 nucleotide spacers separated by 36 nucleotide repeats. The sequences of the repeats were taken from the Csm associated CRISPR cluster of *S*. *thermophilus* DGCC8004. Spacer sequences were exact complements of the targeted region of EGFP, jGFP, or S3 RNA. In the expression cells, the pre-crRNAs are initially cleaved to 72 nucleotide long crRNA, which may be then be trimmed to 40 nucleotide long matured crRNAs with 32 nucleotides of spacer sequence (17).

### Cloning, expression and purification of StCsm complexes

Wt and mutant *Streptococcus thermophilus* Csm (StCsm) complexes were obtained as described previously (17). Briefly, *Escherichia coli* ER2566 (DE3) was transformed with three plasmids: (i) plasmid pCas/Csm which contains a cassette including all the *cas/csm* genes (except *cas1* and *cas2),* (ii) plasmid pCRISPR which contains four identical tandem copies of the repeat-spacer unit flanked by the leader sequence and the terminal repeat, (iii) plasmid pCsm2-Tag which contains a N-terminal-StrepII-tagged variant of *csm2* gene. Next, the ER2566 (DE3) containing these plasmids was grown at 37°C in LB medium supplemented with streptomycin (25 μg/μl), ampicillin (50 μg/μl), and chloramphenicol (30 μg/μl) and expression of wt StCsm complex was induced using 1 mM IPTG. Further, StCsm complex was isolated by subsequent Strep-chelating affinity and size exclusion chromatography steps. The protein composition of the isolated StCsm was analyzed by SDS-PAGE Coomassie staining. StCsm complex containing D33A Csm3 and D16A Cas10 mutants were constructed and isolated as described earlier (17,28). crRNAs co-purified with StCsm were isolated using phenol:chloroform:isoamylalcohol (25:24:1, v/v/v) extraction and precipitated with ethanol. crRNAs were separated on a denaturing 15% polyacrylamide gel (PAAG) and depicted with SybrGold (Thermo Scientific) staining.

### Zebrafish strains and fish maintenance

vasa:EGFP (33) (*Tg(ddx4:ddx4-EGFP*), ZFIN ID: ZDB-TGC0NSTRCT-070814-1) (germline),cmcl2:EGFP (36) (*Tg(myl7:EGFP*), ZFIN ID: ZDB-TGC0NSTRCT-070117-164) (heart), nkx2.5:EGFP (heart) (*Tg(nkx2.5:EGFP*), ZFIN ID: ZDB-TGC0NSTRCT-120828-5) (35) were of ABTL genetic background. The mito:EGFP fish (throughout the embryo) (*Tg(Xla.Eef1a1:mlsEGFP*), ZFIN ID: ZDB-TGC0NSTRCT-090309-1) (34) were of nacre genetic background (38). The fli1:EGFP fish (blood vessels) (*Tg(fli1a:EGFP*), ZFIN ID: ZDB-TGC0NSTRCT-070117-94) (37) were of casper genetic background (39).

The nucleotide sequence of the EGFP transgenes was confirmed by Sanger sequencing in all cases. All reporter lines also contain wild-type copies of the respective genes. General maintenance, collection, and staging of the zebrafish were performed as described previously (40). Embryos were maintained in Danieau zebrafish medium and grown at 28°C. The developmental stages were estimated based on time post-fertilization (hours or days; hpf or dpf) at 28°C.

### Preparation of DNA and RNA substrates

Synthetic oligodeoxynucleotides were purchased from Metabion. All RNA substrates were obtained by *in vitro* transcription using TranscriptAid T7 High Yield Transcription Kit (Thermo Scientific). Briefly, plasmids pSG1154_jGFP, pUC18_S3/1 and pUC18_EGFP were used as a template to produce different DNA fragments by PCR using appropriate primers containing a T7 promoter in front of the desired RNA sequence. RNA substrates were 5’-labeled with [γ^33^P] ATP (Perkin Elemer) and PNK (Thermo Scientific). Ss M13mp18 plasmid DNA was purchased from New England BioLabs. A full description of RNA substrates is provided in Fig. S1.

### Cleavage Assay *in vitro*

The StCsm RNA cleavage reactions *in vitro* were performed at 28°C and contained 8 nM of 5‘-radiolabeled RNA and 160 nM StCsm in the Reaction buffer (33 mM Tris-acetate (pH 7.9 at 25°C), 66 mM K-acetate, 0.1 mg/ml BSA) supplemented with 10mM Mg-acetate. Reactions were initiated by addition of the Mg^2+^. The samples were collected at timed intervals and quenched by mixing 5 μl of reaction mixture with 10 μl of phenol:chloroform:isoamylalcohol (25:24:1, v/v/v). The aqueous phase was collected and mixed with 2x RNA loading buffer (Thermo Scientific) followed by incubation for 7 min at 85°C. The reaction products were separated on a denaturing 15% PAAG and depicted by autoradiography. The StCsm reactions on circular ssDNA *in vitro* were performed at 28°C and contained 1 nM M13mp18 ssDNA, 7.5 nM StCsm, and 7.5 nM RNA in the reaction buffer supplemented with 10 mM MnCl_2_. Reactions were initiated by addition of Mn^2+^. The samples were collected at timed intervals and quenched by mixing 5 μl of reaction mixture with 2x loading dye (98% formaldehyde, 25 mM EDTA, 0.025% bromophenol blue), followed by incubation for 7 min at 85°C. The reaction products were separated during 1% agarose gel electrophoresis in TAE buffer (40 mM Tris, 5 mM Na-acetate, 0.9 mM EDTA, pH 7.9), stained with SYBR Gold (Thermo Scientific) and visualized using Fluorescent Image Analyzer FLA-2000 (Fuji Photo Film, Japan).

### Microinjection

Freshly laid fertilized eggs (or embryos) were collected from breeding tanks and injected with 1 nl of 0.5mg/ml StCsm complex into the yolk of one cell stage embryos using an Eppendorf FemtoJet microinjection setup. The embryos were incubated afterwards at 28 °C.

### Microscopy

Embryos were anaesthetized using tricaine. Fluorescence from live embryos was observed using a Leica M165FC fluorescent microscope equipped with a Leica DFC450C digital camera.

### Flow cytometry analysis

About 20-30 embryos at 48 hpf were collected randomly and washed twice with Hank’s solution. The embryos were minced in a Petri dish with a fine scalpel. 800 μl of 0.25 % trypsin were added and mixed at 500 rpm on a bench top shaker for 1h at RT. The cells were filtered through a 40 μm cell strainer and washed 3 times with Hank’s solution. The remaining was centrifuged at 2000 rpm. The pellet was resuspended in 500 μl Hank’s solution and analyzed for EGFP fluorescence by BD FACS Calibur.

## RESULTS

### Study design

We have tested whether Csm complexes can be used for mRNA knockdown in zebrafish. In order to compare efficacies in different tissues and in different developmental stages in a meaningful way, we did not target endogenous messenger RNAs. Instead, we targeted the EGFP transcript expressed as a transgene from a variety of different promoters. In order to judge the specificity of the mRNA targeting, we used not only a crRNA guide that was optimal for the EGFP transcript in zebrafish, but also a crRNA guide that was optimized for the jellyfish GFP (jGFP) transcript and annealed to the EGFP transcript with three clustered and three separate mismatches, and a completely unrelated S3 crRNA guide (Fig. S1). In order to distinguish between Csm RNase and DNase dependent effects, we also prepared variants of the crRNA Csm complexes that differed from the wild-type complex by the D33A substitution in Csm3 (DNase activity only) and the D16A substitution in Cas10 (RNase activity only). Complexes were injected into the yolk of zebrafish at the 1-cell stage, and EGFP fluorescence was then monitored at different time points, either by direct observation or FACS analysis of trypsin-digested embryos (Fig. 1).

**Figure 1.**
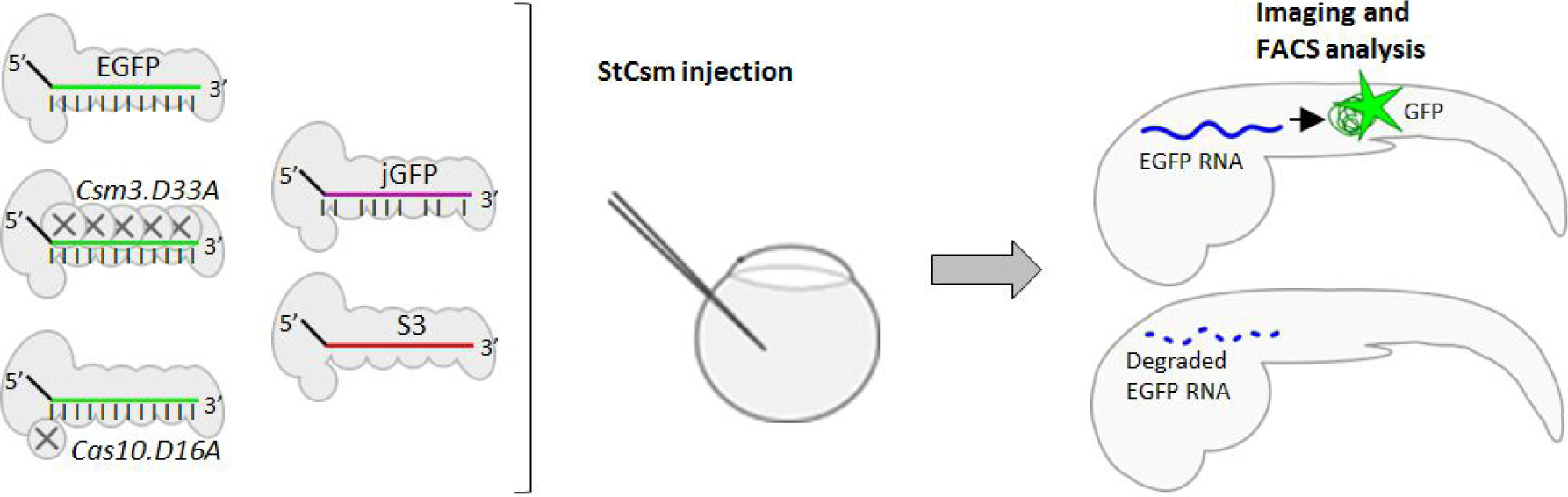
Experimental design: Purified wild-type, DNase (D16A) or RNase (D33A) defective variants of StCsm complexes targeted to degrade EGFP transcript were injected into the yolk of zebrafish embryos. Suppression of EGFP fluorescence was monitored by imaging and FACS analysis. Level of crRNA complementarity to EGFP transcript is indicated by vertical bars. See also Fig. S1.

### Bacterial expression and purification of StCsm-crRNA complexes

In order to generate pre-crRNAs for loading of Csm complexes in the *Escherichia coli* expression host, synthetic CRISPRs were generated. These contained the 36 nucleotide repeats found in the Cas/Csm associated CRISPR cluster of S. *thermophilus* DGCC8004, and four identical spacers of reverse complementary sequence to the targeted region of EGFP, jGFP or S3 RNA. The pre-crRNAs were then co-expressed with the S. *thermophilus* Csm genes and co-purified with processed crRNA. Moreover, the Cas10 D16A and Csm3 D33A variants of the StCsm complex were also prepared together with EGFP crRNA (17). The purified Csm(EGFP), Csm(jGFP), Csm(S3), D16A Csm(EGFP) and D33A Csm(EGFP) complexes were checked for protein purity by denaturing gel electrophoresis and Coomassie staining (Fig. 2A). Their crRNA content was analyzed by denaturing PAGE analysis followed by RNA SYBR Gold staining (Figs. 2A and 2B). The results confirm essentially equal protein content of all complexes, and approximately equal crRNA content of all complexes. Csm complexes contained a mixture of a 72 nucleotide long crRNA arising from cleavage of the pre-crRNA, and of 40 nucleotide long crRNA arising from further trimming of this crRNA, as described earlier (17).

**Figure 2.**
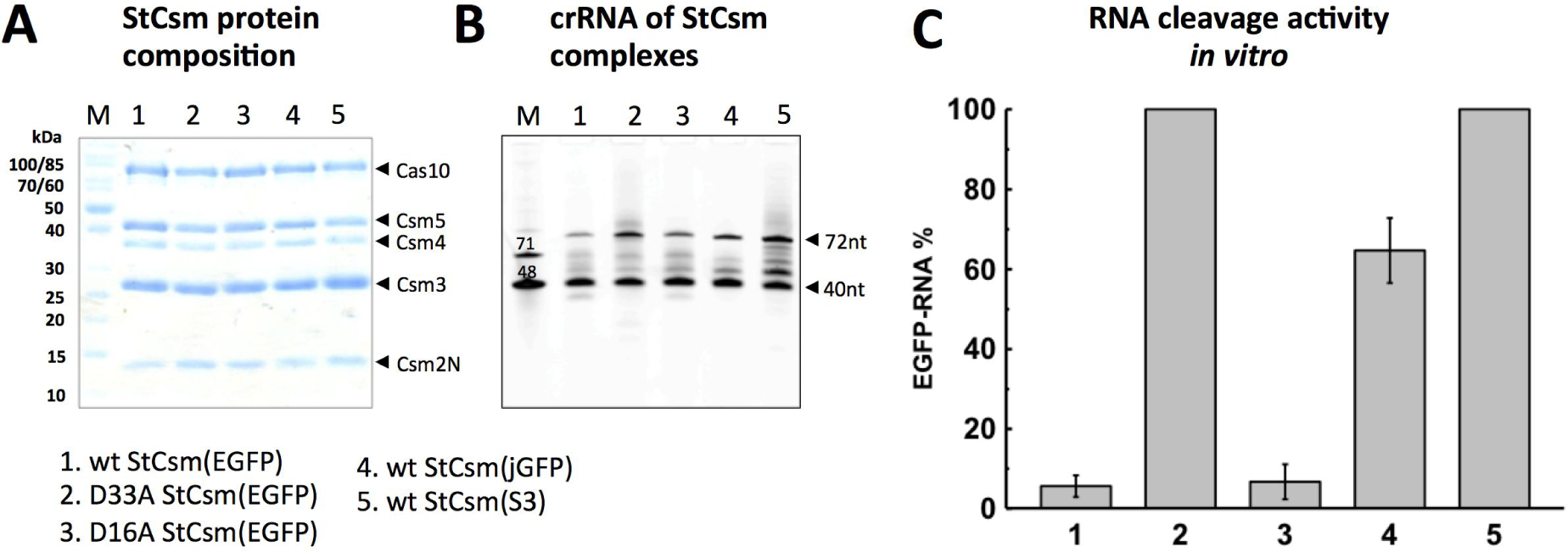
*In vitro* characterization of StCsm complexes: (A) Coomassie blue-stained SDS-PAGE of purified StCsm complexes. M, protein mass marker. (B) Denaturing PAGE analysis of crRNA co-purifying with the StCsm complexes. M, synthetic DNA marker. (C) RNA cleavage activity *in vitro.* Depletion of radioactively labeled target EGFP RNA measured in 1 hour after RNA cleavage reaction initiation. Data are represented as mean ± SEM. See also Figs. S2, S3.

### Test tube validation of the RNase and DNase activities of Csm-crRNA complexes

The RNase activity of the StCsm complexes was tested at 28 °C, the temperature used for the zebrafish experiments. Cleavage of a radiolabelled RNA fragment containing the EGFP target region was monitored over time (Fig. S2), and the remaining amount of substrate was quantified after one hour of incubation (Fig. 2C). The wt StCsm(EGFP) cleaved complementary substrate RNA with the characteristic 6-nucleotide stagger described earlier (17,32). In contrast, the amount of substrate remained essentially unchanged after the 1h incubation period with the RNase defective D33A variant. In contrast, the DNase defective D16A mutant cleaved the substrate RNA like the wild-type StCsm complex. Mismatches between guide and target sequence hindered RNA cleavage more effectively at lower than at higher temperature (data not shown). At 28 °C, incubation of the substrate with StCsm(jGFP) (three clustered and three isolated mismatches between crRNA and target) left most of the substrate undigested, arguing for good specificity of the StCsm crRNA complexes. Incubation of substrates with an irrelevant (S3) guide did not cause any RNA cleavage (Fig. 2 and Fig. S2).

The DNase activity of the StCsm complexes was tested with a single stranded DNA substrate in the presence of target RNA mimicking a transcript, again at 28 °C, the relevant temperature for the zebrafish experiments. In agreement with earlier data (28), robust DNA degradation was observed when StCsm crRNA complexes were used together with matched RNAs. However, DNA degradation appeared to occur on a slower timescale than RNA degradation. As predicted, DNA degradation unaffected by the D33A mutation, but completely impaired by the D16A mutation. The StCsm(jGFP) and StCsm(S3) complexes that had mismatched and unrelated guides to the EGFP RNA did not cleave DNA, even though they were active in the presence of their matched RNAs (Fig. S3). We conclude that RNA dependent DNA degradation is also very specific.

### Knockdown of the maternal vasa:EGFP

The fluorescence of vasa:EGFP fish in our hands was consistent with literature reports (33,41). We did not observe EGFP fluorescence up to 5 dpf (the last time point checked due to legal restrictions) embryos that were derived from crosses of wt females with vasa:EGFP males. This observation confirms that vasa:EGFP is exclusively expressed from the maternal transcript and that transcription of the EGFP transgene-unlike the transcription of endogenous vasa - does not set in at the gastrulation stage (Fig. S4).

In initial experiments, we determined a minimal dose of StCsm(EGFP) to achieve knockdown (Fig. S5). In the following, we describe only experiments with this optimal dose. When we injected 1 nl of 0. 5 mg/ml StCsm(EGFP) crRNA into the yolk of 1-cell stage vasa:EGFP embryos (from mating’s of vasa:EGFP mothers and fathers), we did not observe differences in fluorescence between non-injected embryos, mock-injected embryos, or embryos injected with the StCsm, in the early stages of development (3 hpf), presumably due to maternally deposited protein. However, fluorescent imaging of the embryos at 1 dpf to 5 dpf showed clear reduction of fluorescence in the germ cells of embryos injected with wt StCsm(EGFP) compared to non-injected controls. Interestingly, fluorescence from these cells (above background in the entire embryo) was almost completely abolished at 1 dpf (Fig. 3). However, in a small fraction of fish, the injections did not have a clear-cut effect, suggesting technical issues rather than a knockdown failure (no correction for injection failure was made, and all fish for which injection was attempted were included in the subsequent quantitative analysis). The loss of fluorescence compared to control in the germ cells of wt StCsm(EGFP) injected embryos was persistent at least during the first five days of development. Knockdown did not affect the (weak) background fluorescence initially in the entire embryo and primarily in the brain, which may be explained by maternal deposition of EGFP protein.

**Figure 3.**
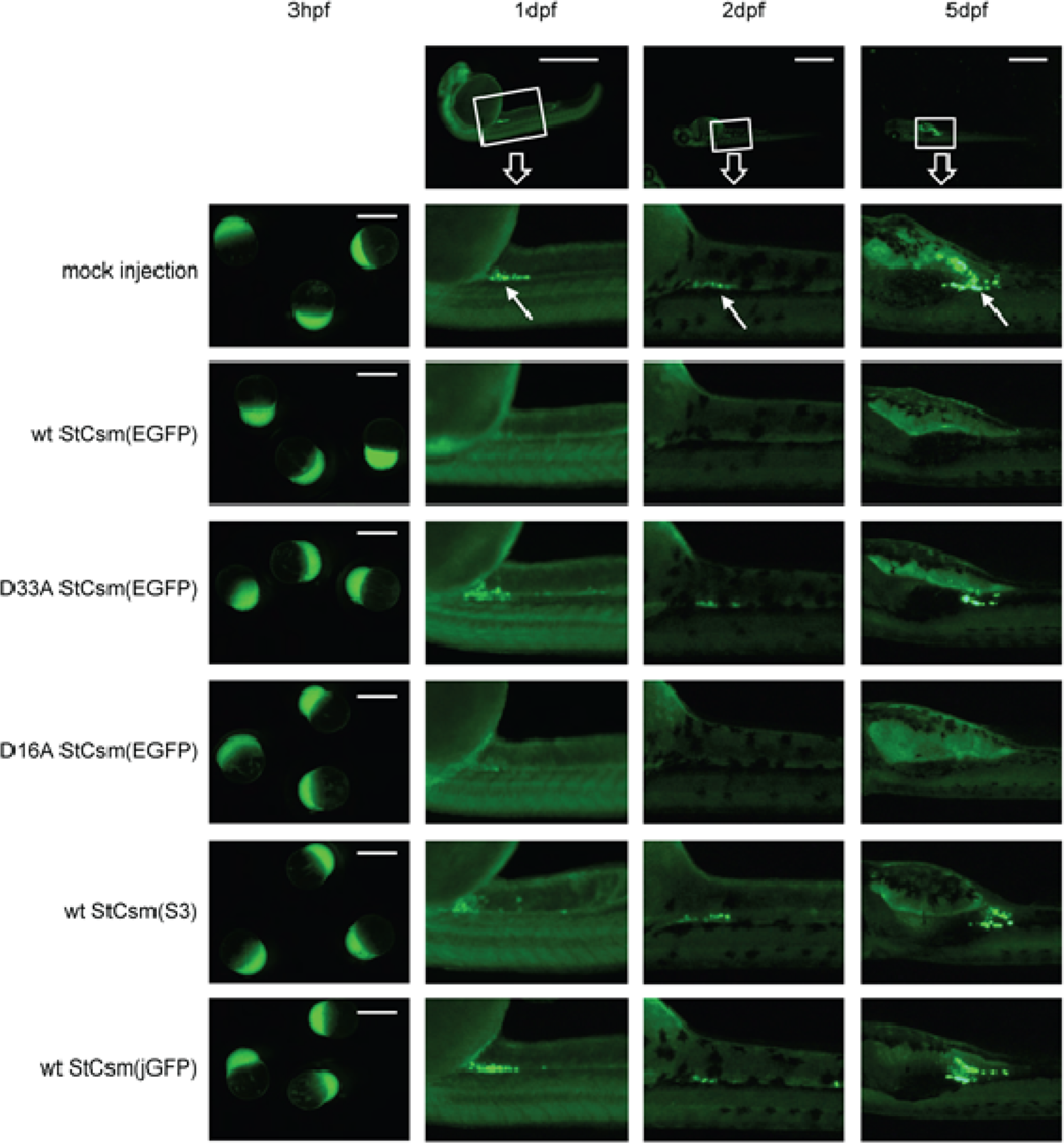
Microsopy of StCsm mediated EGFP knockdown in vasa:EGFP fish: Fluorescence from vasa:EGFP fish after injection with wt or mutant St-Csm-crRNA complexes. The arrows indicate the location of primordial germ cells. Injection was done at the 1-cell stage, observation were 3 hpf, 1 dpf, 2 dpf and 5 dpf. The scale bar represents 1 mm. See also Figs. S4, S5.

As the EGFP in the vasa:EGFP strain is expressed exclusively from the maternal RNA, the loss of fluorescence should be the result of RNA knockdown, and should not be dependent on potential effects of the DNase activity of the complex on the transgene itself. In order to verify this prediction biochemically, we injected embryos with the DNase dead mutant D16A StCsm(EGFP). The D16A StCsm(EGFP) extinguished fluorescence at 1 dpf like the wild-type protein. As a further control, we injected embryos instead with the RNase dead D33A variant of the Csm-crRNA complex. The mutant did not cause a noticeable decrease in fluorescence, confirming that the decrease was indeed due to RNA knockdown and not to indirect effects on RNA. Finally, we checked the consequences of imperfect reverse complementarity between jGFP crRNA and EGFP mRNA. The six mismatches between the two RNAs appeared to abolish EGFP degradation (Fig. 3).

In order to make the above observations quantitative, we minced and trypsinized embryos from mating of a single pair of fish and subjected the resulting pool of cells (and aggregates of cells) to fluorescence activated cell sorting. In order to avoid the influence of background fluorescence, the gating window in the FACS was set to count cells only in the region of very high fluorescence (50-fold above mean). The number of cells with fluorescence above the threshold was then used as a measure of fluorescence. Analysis of 1 dpf and 2 dpf embryos showed 20-fold and 6-fold reductions respectively in the numbers cells with high EGFP fluorescence with wt and D16A StCsm(EGFP). No reduction of fluorescence was observed using the RNase dead D33A StCsm(EGFP), the wt StCsm(jGFP) complex with imperfectly complementary RNA or the control crRNA wt StCsm(S3) (Fig. 4A). Differences were significant independently of the precise choice of threshold for highly fluorescent cells (not shown). The highly fluorescent cells showed a high side scatter which is consistent with the increased size of primordial germ cells (42).

**Figure 4.**
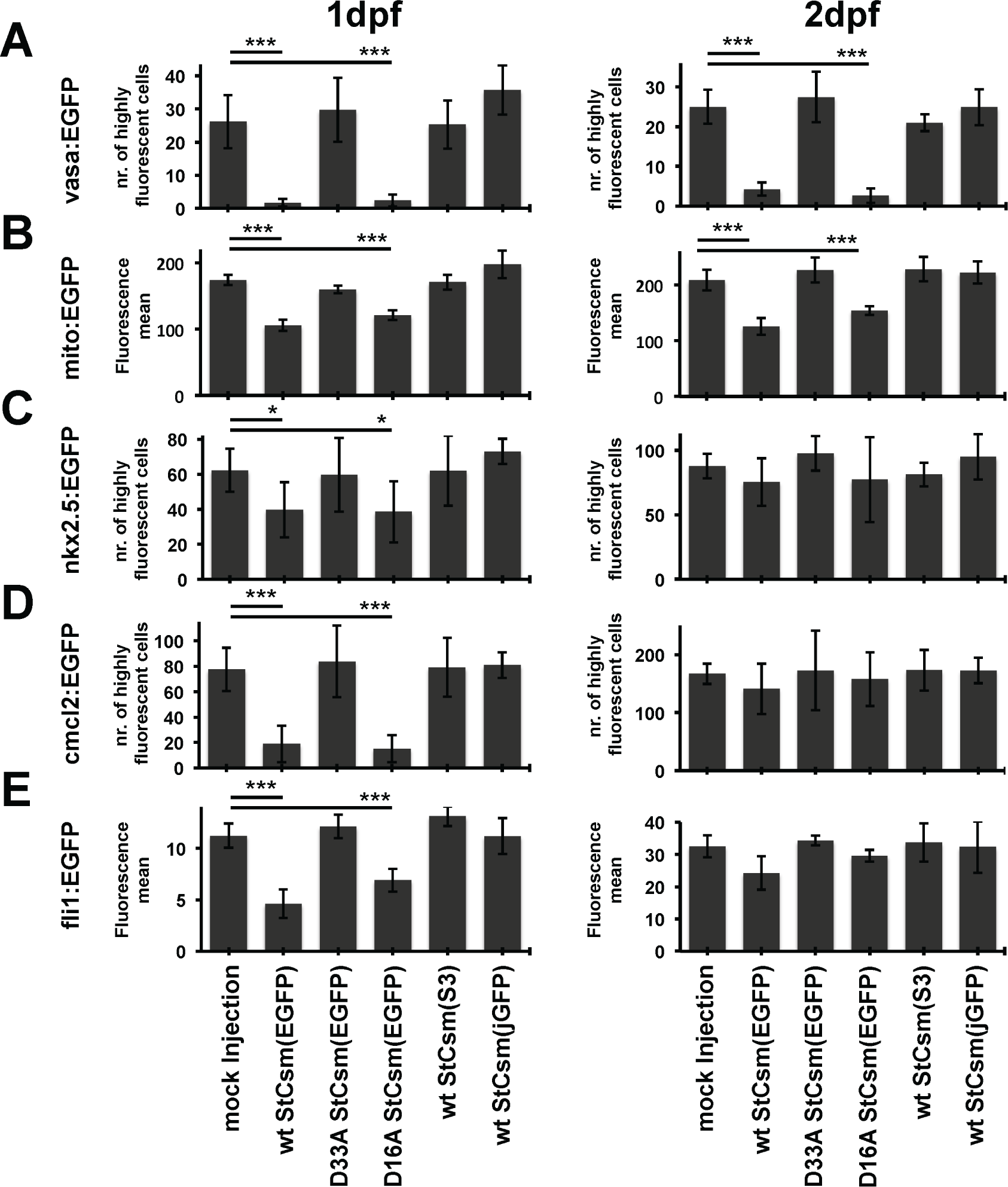
Quantification of knockdown efficiency by FACS analysis: EGFP fluorescence in (A) vasa:EGFP, (B) mito:EGFP, (C) nkx2.5:EGFP, (D) cmcl2:EGFP and (E) fli1:EGFP fish was quantified 1 dpf and 2 dpf by flow cytometry of minced and trypsinized embryos. For vasa:EGFP, nkx2.5:EGFP cmcl2:EGFP embryos, the number of highly fluorescent cells (at least 50 times background fluorescence was counted). Results are from three independent experiments, error bars represent one standard deviation. * P < 0.05 as compared with respective controls, *** P < 0.001 as compared with respective controls. Data are represented as mean ± SEM. See also Figs. S6-S15.

### Knockdown of the mixed maternal-zygotic mito:EGFP

Embryos from mito:EGFP show green fluorescence in mitochondria already in oocytes, and
throughout the life of the fish. In crosses between wild-type ABTL fish and mito:EGFP fish,fluorescence is observed throughout embryonic development when the mother carries the mito:EGFP transgene, indicating maternal and then zygotic expression, whereas fluorescence is observed only after the midblastula transition in the reciprocal cross (Fig. S6). For knockdown experiments, we used the same concentrations of StCsm(EGFP) and variants as in the vasa:EGFP experiment. The lines were analyzed by fluorescence microscopy and FACS analysis of digested embryos at 1 dpf, and in the case of the mito:EGFP x mito:EGPF crosses, also at 2 dpf. As green fluorescent is present throughput the embryo and not concentrated in a particular cell type, we quantified mean fluorescence, instead of the number of highly fluorescent cells as for the vasa:EGFP fish.

When we used fish from mito:EGFP mothers and wild-type fathers, knockdown efficiency using wild-type StCsm(EGFP) was quite good. The knockdown efficiency was insignificantly lower with the D16A variant, arguing for a possible contribution of DNA cleavage in the observed reduction of fluorescence. However, the D33A variant, which has only DNase activity, was completely ineffective, suggesting that the reduction of fluorescence was not due StCsm(EGFP) DNase activity (Fig. S7A). When we repeated the experiment for the reciprocal cross, the reduction in fluorescence that could be achieved by StCsm(EGFP) injection was much lower, either because maternal RNA overwhelms StCsm(EGFP) capacity, or more likely, because fluorescence of maternally deposited EGFP protein is not affected by the knockdown (Fig. S7B). A similarly low knockdown efficiency was also observed when the StCsm(EGFP) was pitched against maternally deposited RNA and zygotically expressed RNA in embryos from crosses of mito:EGFP parents (Fig. 4B4 and Fig. S8).

### Knockdown of the zygotic cmcl2:EGFP and fli1:EGFP, but not nkx2.5:EGFP

In order to confirm that StCsm(EGFP) could also be used to for knockdown of zygotically expressed mRNAs, we tested knockdown of EGFP transcripts from other promoters. Consistent with predominantly zygotic promoter activity, heterozygotic nkx2.5:EGFP, cmcl2:EGFP or fli1:EGFP fish carrying a maternally or paternally inherited transgene exhibited similar fluorescence at all tested time points (4 hpf, 1 dpf, 2 dpf, 5 dpf) (Fig. S9-S11). As before, the amount of StCsm(EGFP) for injections was not optimized anew and the same concentration as for the vasa:EGFP experiments was used throughout. All lines were analyzed by fluorescence microscopy and FACS analysis of digested embryos at 1 dpf and 2 dpf.

The nkx2.5:EGFP and cmcl2:EGFP fish exhibit fluorescence in the heart. We quantified knockdown efficiency as for the vasa:EGFP line by counting the number of highly fluorescent cells (50-fold more fluorescent than background). A knockdown of 30-40% was detected in the nkx2.5:EGFP fish upon StCsm(EGFP) injection by FACS (Fig. 4C). This reduction in EGFP fluorescence was not detectable by imaging. (Fig. S12). A stronger, four-fold fluorescence reduction could be achieved in the cmcl2:EGFP line at 1 dpf (Fig. 4D and Fig. S13). As the controls with the D16A and D33 Csm(EGFP) variants showed, the reduction of fluorescence was almost exclusively due to the RNase activity of the StCsm(EGFP) complex in both cases. At 2 dpf, the effect of knockdown had essentially faded away.

The fli:EGFP fish exhibit fluorescent vasculature throughout the embryo. Therefore, we quantified knockdown efficacy by mean fluorescence in this case. At 1 dpf, more than 50% knockdown was achieved using the wt StCsm(EGFP) complex. The reduction of EGFP fluorescence was slightly lower with the D16A variant, raising the possibility that the DNase activity of the Csm complex may have contributed the reduction of fluorescence. However, the RNase deficient version of the complex did was ineffective. At 2 dpf, knockdown effects were reduced to statistically insignificant levels (Fig. 4E and Fig. S14).

### Low StCsm(EGFP) toxicity

StCsm(EGFP) injection was not obviously toxic. Some developmental defects of injected fish were observed in the observation period until 5dpf, but the rate was similar for mock- and StCsm(EGFP) injected embryos and did not increase even when the amount of StCsm(EGFP) was fivefold higher than in knockdown experiments. In this respect, StCsm crRNA complexes may be superior to morpholinos (Fig. S15).

## DISCUSSION

### Csm mediated reduction of EGFP fluorescence is due to the Csm RNase activity

Experiments with Csm variants show that reductions in EGFP fluorescence are largely independent of the DNase activity of the Csm complex, even for EGFP expressed from zygotically active promoters. This could reflect faster RNA than DNA degradation (in part due to transcriptional silence up to the maternal to zygotic transition (MZT)), or the exclusion of the StCsm complex from the nucleus due its large size and the absence of a nuclear localization signal (NLS). It remains to be tested how nuclear targeting of mRNAs or pre-mRNAs by StCsms with NLS compares to cytoplasmic targeting. Although we did not see evidence for DNase activity of the Csm complex in our experiments, we recommend using the “cleaner” Csm D16A variant for RNA knockdown experiments.

### Csm mediated knockdown is best for maternally deposited RNAs

Csm mediated RNA knockdown worked best for the exclusively maternal EGFP transcript generated under the control of the vasa promoter. The knockdown for transcripts that are expressed in the zygote was less pronounced than for the maternally deposited RNA, and the effect decreased over time. Observation of significant knockdown effects for the cmcl2 and fli1 promoter driven EGFP transcripts strongly suggests that the Csm bound crRNA survives the widespread RNA degradation during the MZT at 3.5 hpf, most likely due to protection of the crRNA along its entire length by the Csm complex (17). At 1 dpf, EGFP fluorescence was reduced less than two-fold in the mito:EGFP and nkx2.5:EGFP lines. As most vertebrate genes are haplo-sufficient (43,44), this efficiency may not be enough to elicit phenotypes. Therefore, for now we recommend Csm mediated knockdown only for maternally deposited RNAs and for use very early in development (at most 2dpf).

### Comparison to other knockdown methods

With maternal RNA transcripts within its scope, Csm knockdown currently fills a niche for knockdown in zebrafish that is not well addressed by the morpholino approach and inaccessible to the dCas9 transcription suppression approach. As zygotic transcription sets in late in zebrafish, many important biological questions, including questions of cell fate, RNA translation control, and others, can be addressed during this time window. For programmable enzymatic degradation of transcripts, NgAgo is currently the only competitor to the Csms that has been tested in zebrafish. The data do not suffice to address the relative merits of the two approaches. The NgAgo targeted *fabp11a* transcript is expressed as early as the zygotically expressed EGFPs in this study, making it hard to compare knockdown persistency. NgAgo has the advantage over Csms that it can be loaded in a test tube with guide nucleic acid. However, with its DNA guide may endanger genome stability more than a DNase deficient Csm complex with its RNA guide. NgAgo cuts RNA at a single site whereas Csm complexes make multiple cuts in a target RNA. Whether this difference matters in the background of a host such as zebrafish that has pathways for the degradation of uncapped or non-polyadenylated mRNAs remains to be explored.

### Outlook

In this work, we have demonstrated the utility of StCsm for programmable degradation of maternally deposited mRNA in zebrafish. Exclusion from the nucleus effectively attenuated or abolished the DNase activity of the Csm complex. Nevertheless, we recommend the “cleaner” StCsm D16A variant for targeted knockdown experiments. We see the current demonstration as “proof of principle”, but expect that the method can be improved. We have so far only tested the Csm from one bacterial species, and even with this Csm complex, there may be room for improvement. It is possible that targeting pre-mRNAs and RNAs at the source in the nucleus (using a Csm complex with NLS) could be more effective than cytoplasmic targeting for zygotic transcripts. Moreover, we have only used preformed StCsm(EGFP) complexes and not attempted to express and assemble StCsm(EGFP) complexes from injected RNAs or RNAs made by transcription in zebrafish from DNA templates. We have only injected into the yolk and not the embryo proper, and we have not explored the dose dependence of the knockdown efficiency. We have also not attempted to stabilize the crRNA or Csm proteins against degradation, and most importantly, we have not tested other CRISPR endoribonucleases, which may be easier to load with guide nucleic acids.

## SUPPLEMENTARY DATA

Supplementary Data are available at NAR online.

## ACKNOWLEDGEMENT

We thank the labs of Lisbeth Charlotte Olsen, Huai-Jen Tsai, Gergana Dobreva, Seok-Yong Choi and Brant M.Weinstein who created the EGFP zebrafish lines used in this work, and Prof. Jacek Kuznicki, Prof. Agnieszka Chacinska, Dr. Cecilia Winata (all IIMCB) for passing them on to us. Dr. Małgorzata Wiweger, Dr. Lidia Wolińska-Nizioł and the staff of the IIMCB zebrafish facility are gratefully acknowledged for help with animal husbandry. We thank Mantvyda M. Grusyte for help with plasmid construction. We thank Dr. Lina Jakutyte-Giraitiene for the kind gift of pSG1154_jGFP.

